# Rapid evolution in response to biological invasion: variation in resistance and survival of the European bitterling fish across a gradient of *Sinanodonta woodiana* expansion

**DOI:** 10.1101/2025.09.02.673702

**Authors:** Abhishek Nair Anil, Dariusz Halabowski, Kacper Pyrzanowski, Grzegorz Zięba, Joanna Grabowska, Carl Smith, Karel Douda, Martin Reichard

## Abstract

Biological invasions impose new selective pressures on native species, prompting rapid evolutionary responses. We examined how the parasitic larvae (glochidia) of the invasive East Asian mussel *Sinanodonta woodiana* affect European bitterling fish (*Rhodeus amarus*) populations with different histories of exposure to the parasite (none, 15 years, or 40+ years). Glochidia parasitise fish fins and gills, and bitterling are especially vulnerable to parasitism due to their spawning association with mussels. We found the lowest glochidia load in bitterling from populations with the longest association with *S. woodiana*. While infection reduced swimming capacity proportionally to parasite load, this effect was not population-specific. However, post-infection mortality was highest in naïve and lowest in long-associated populations, indicating evolved tolerance. No long-term effects on survival, growth, condition, or reproduction were observed, suggesting fitness costs are concentrated shortly after infection. These results show that novel host–parasite interactions during invasions can drive rapid adaptive change, emphasising the need to consider evolutionary dynamics in invasion management.

## INTRODUCTION

Biological invasions increasingly represent one of the dominant drivers of natural selection. As anthropogenic activity facilitates transport of organisms to new geographic areas, new species enter ecological interactions with native organisms. This situation produces novel selective pressures, both direct (leading to coevolution) and indirect (leading to adaptation to new ecological and environmental conditions). Non-native species typically lack their original partners in interspecific positive (e.g. pollination, dispersal) and negative (competition, parasitism) associations. This absence may hinder or prevent their establishment in the new region (Richardson et al. 2000; Mitchell and Power 2003) or provide benefits through the vulnerability of native species arising from their evolutionary naivety, exacerbating the detrimental impacts of invaders on native species and communities (Cox 2004; Cox and Lima 2006; Sih et al. 2010).

Relationships between native and invasive species are dynamic. While rapid change may arise from experiences at the behavioural level (Cox and Lima 2006), such as learning the unpalatability of a potentially novel prey (Berger et al. 2001), many responses are evolutionary. For example, interaction with a non-native parasite can cause rapid evolutionary responses in the host, leading to mitigation of the impacts of new parasitic species (Decaestecker et al. 2007; Morran et al. 2011). Hosts respond to parasites through the evolution of resistance (to decrease parasitic burden) or tolerance (to optimise the energetic cost of a parasitic load) (Råberg 2009; Svensson and Råberg 2010). Defence against parasites is energetically costly (Sheldon and Verhulst 1996) and maintaining active resistance increases host metabolic rate (Martin et al. 2002), while tolerance requires higher energetic uptake by the host (Knutie et al. 2016). Hence, both strategies are energetically costly (Thambithurai et al. 2022) and are traded off against other energy-demanding organismal functions. These are not exclusive outcomes, and multiple strategies can be deployed to mitigate parasite load. For example, Knutie et al. (2017) showed that the Cuban tree frog exhibits tolerance and resistance against a gut nematode parasite. In this study, frogs simultaneously limited infection intensity (resistance) and maintained growth and body condition despite parasite burden (tolerance), showing that both strategies can act in parallel within individuals to reduce the negative effects of parasitism. The two mechanisms of coping with a parasite can also influence how natural selection maintains the variation in defence capabilities of host individuals within a population (Parker et al. 2014) and produce parasite-mediated selection (Coltman et al. 1999).

Ectoparasites impose immediate impacts on fish hosts, such as breakage of epithelial tissues, hydromineral imbalances and disturbances to metabolism (Douda et al. 2017a), and resisting these impacts is energetically expensive. Short-term effects are often seen as departures from standard behaviour, impairment in physiological capacity such as swimming performance or alteration of ventilation rates (Martin et al. 2002; Husak et al. 2016; Slavík et al. 2017). They may also lead to major impacts, including mortality. Longer-term costs of parasites are often observed as an energetic trade-off between coping with a parasitic load and other physiological demands on host individuals. Slower growth rate, reduced reproductive condition and increased mortality are often reported as long-term effects of parasitism (French et al. 2009; Barber et al. 2011; Taeubert and Geist 2013). Parasites often initiate host stress responses mediated by release of stress hormones, such as increases in plasma cortisol (Douda et al. 2018; Reichard et al. 2023). Indirect effects of parasite infection mediated via chronic stress can also produce long-lasting impacts on survival, growth and reproductive output (Saarinen and Taskinen 2005; Neuman-Lee et al. 2015; Seguel et al. 2019).

Rapid coevolution refers to evolutionary changes to adapt to interacting species at an unusually rapid rate, usually within several generations. Antagonistic interactions are key drivers for rapid adaptive evolution and can influence pathogen-mediated fitness costs (Marston et al. 2012). Hosts may evolve rapidly to reduce the fitness costs imposed by novel parasites, resulting in fine-scale variation in defence traits and population-level differentiation. Biological invasions introduce new selective regimes by bringing evolutionary naïve hosts and parasites together. These novel interactions provide powerful natural experiments for studying the mechanisms and outcomes of rapid coevolution, with broad implications for understanding the evolutionary basis of host defence, community resilience, and species persistence under environmental change (Facon et al. 2006; Hufbauer et al. 2012). One particularly relevant mechanism is selective disappearance, in which individuals that fail to cope with parasite-induced stress or damage experience higher mortality. Such mortality may drive strong selection favouring resistant or tolerant host phenotypes, making selective disappearance a potentially important force in parasite-mediated rapid evolution (Sheldon and Verhulst 1996).

Here, we use the relationship between European bitterling fish (*Rhodeus amarus*) and invasive freshwater mussel (*Sinanodonta woodiana*) to test rapid host responses to a novel parasitic species. The European bitterling is a small cyprinid fish widespread in Europe (Smith et al. 2004). Geographically, bitterling diversity is highest in East Asia with only a single species group present in West Palearctic (Chang et al. 2014; Bartáková et al. 2019). Bitterling and unionid mussels live in an unusual reciprocal parasitic relationship. Female bitterling deposit their eggs in the gills of mussels, where they are protected for several weeks and interfere with mussel feeding and gas exchange. In turn, parasitic larvae of unionid mussels obligately attach to fish, possibly including the bitterling, before metamorphosing to juvenile mussels. *R. amarus* has coevolved with several species of European mussels and can use them as hosts for their eggs. European bitterling also resist the glochidia of European unionid mussels compared to other potential hosts among European fishes (Smith et al. 2004; Blažek and Gelnar 2006; Reichard et al. 2010; Huber and Geist 2019).

The invasive mussel *Sinanodonta woodiana* occurs naturally in East Asia and has invaded European freshwaters within the last 60 years (Konečný et al. 2018). Its glochidia can use a wide range of native species in East Asia, and such generalist host use has been implicated in its successful invasion of Europe and elsewhere (Douda et al. 2012). Invasive *S. woodiana* has reversed the bitterling-mussel association in Europe. Unlike European mussels, it does not host European bitterling eggs (Smith et al. 2004; Marčić et al. 2024; Halabowski et al. 2025). At the same time, it effectively uses *R. amarus* as a host for its glochidia (Douda et al. 2017b). This contrast with native European mussels has been ascribed to contrasting selection from multiple bitterling species on *S. woodiana*’s ability to resist bitterling oviposition, while *R. amarus* in Europe has experienced low resistance to infection by glochidia of European mussels (Reichard et al. 2006, 2010). At the glochidial stage of their association, strong local adaptation of bitterling hosts, as well as strong selection in the coevolutionary hotspot of the bitterling-mussel interaction in East Asia, produce competent immune responses toward glochidia of native but not invasive mussel species (Anil et al. 2024). In Europe, *S. woodiana* initially invaded artificially heated lakes (Kraszewski and Zdanowski 2007) but in the last 20 years started to spread to water bodies with more typical European thermal regimes (Douda et al. 2024; Mehler et al. 2024). This staged expansion creates a gradient of coexistence between *S. woodiana* and *R. amarus* populations and provides an opportunity to test how host populations have evolved to respond to the new parasite across the 40 years of its invasion.

We tested immediate and long-term effects of *S. woodiana* glochidia on bitterling condition and survival. We experimentally infected juvenile fish and subjected them to swimming performance tests. We determined glochidia load and initial survival rates of fish for a period of two weeks post-infection. For long-term effects, we monitored fish condition for six months after parasite attachment, including an overwintering phase. Mortality, body size, and condition factor were recorded to estimate long-term survival and growth rates. At the end of the overwintering period, gonadosomatic index (GSI) and number of oocytes were estimated to test the effect of parasitic load on reproductive allotment. Bitterlings used in the study originated from three populations along a temporal gradient of their historical interaction with *S. woodiana* in Poland, where the mussel has been present since the 1980s (Kraszewski and Zdanowski 2007). We used these populations: old (coexistence of at least 40 years), intermediate (15 years) and without any contact with *S. woodiana,* to test for the role of rapid evolutionary response to mitigate costs of glochidia parasitism.

We predicted a decrease in overall fish condition in the infected group compared to the control group. We also predicted lower glochidia load in fish from populations with longer coexistence with the parasite, in the case of rapid evolution of resistance, but no interpopulation difference in initial glochidia load if evolutionary response is mainly toward increased tolerance. During the initial phase of infection, the number of glochidia attached to individuals is usually high, causing an acute stress response in the host and thus a significant impact on host physiology (Douda et al. 2018; Reichard et al. 2023). As glochidia are known to impose an energetic cost on host fish, we expected attached glochidia to hinder the swimming performance and survival of juvenile fish immediately after infection. We also hypothesised a negative long-term effect of parasite load on the physiology and reproductive traits of the infected fish, demonstrated in a decrease in survival, growth rate and reproductive allotment caused by a combination of parasite load and the challenging overwintering period.

We hypothesised a positive correlation, at the population level, between longer coexistence with the invasive *S. woodiana* and the ability to reduce the cost of parasitism in all tested parameters. Such a result would support the hypothesis of rapid evolution of host response to a non-native parasite. To disentangle the nature of rapid evolutionary response, we compared glochidia load across the gradient of association. Reduction of parasite load in the old-association population (followed by intermediate-association population) would support rapid evolution of resistance mechanisms. Alternatively, no interpopulation differences in parasite load but a decrease in (long-term) cost of parasitism with increasing coevolutionary history would support the hypothesis of the rapid evolution of tolerance in the bitterling.

## METHODS

### Study animals and experimental design

The study used three populations of *R. amarus* from different regions of Poland. Populations of fish were chosen based on the time of the first reports of *S. woodiana* at specific sites according to a published database (Mehler et al. 2024), supplemented by personal communication with A. M. Łabęcka, the source of data for the Mehler et al. (2024) database. We used one population from a site without invasive *S. woodiana* and two populations with contrasting durations of exposure to *S. woodiana*. (1) ‘Without’ – population naïve to *S. woodiana* (River Drzewiczka; N 51.45037, E 20.48654), (2) ‘Intermediate’ - *S. woodiana* first recorded in 2010 (Krajskie Oxbow Lake; N 50.012960, E 19.530882), (3) ‘Old’- *S. woodiana* first recorded in the early 1980s (Lake Licheńskie; N 52.312995, E 18.349566). Fish used in the experiment were young of the year juveniles which were born in captivity to wild-captured parents. All juvenile fish were housed outdoors in fibreglass tanks (1.3 ×□1.3 ×□1.0□m) before the start of the experiment.

For experimental infection with *S. woodiana*, we followed the protocol of Douda et al. (2012). On 28 August 2023, *S. woodiana* were collected from an independent site (not sympatric with any study bitterling populations). Three female mussels were used to extract glochidia, which were pooled in a tank filled with aged tap water and kept in homogenous suspension by continuous mixing (mean suspension density was 4067 viable glochidia per litre). The suspension was divided into six larger and three smaller containers (20 ×□30 ×□10□cm with 30-40 fish in each container for the overwintering experiment and 12 × 15 × 10 cm with 12 fish in each container for the swim tunnel experiment) to infect treatment fish. Water temperature in the infection tanks was 21–22°C. After 10□min of inoculation, fish were transferred into an aerated bath of aged fresh water for 10□min to rinse off any unattached glochidia. Control fish were processed through the same protocol, but with clean water instead of a glochidia suspension.

### Swim tunnel experiment

A subset of experimentally infected fish from all three populations (n=11-12 per population), along with an uninfected group of fish as control, were used to test the role of glochidia infection on swimming performance. A Loligo^®^ mini swim tunnel (Blazka-type, 1500 mL) was used to test the effect of parasite load on critical swimming speed (Ucrit) of individual fish. All swim tunnel runs were completed within four days after experimental infection (while fish possessed an initial glochidia load). The fish were fed with chopped bloodworm until 24 hours before the swimming performance test.

Water velocity in the swim tunnel at different velocity settings was calibrated by the tracking method using food dye as suggested by the manufacturer. Experimental runs were conducted with the tunnel motor initially set to 0.6 BL/s (BL: body length of individual fish) with a constant velocity increment of 0.8 BL/s that increased every 150 seconds. Motor settings for swimming performance trials were determined by a pilot study with *R. amarus* juveniles to attain water velocities sufficient for bitterling juveniles to reach exhaustion.

Juvenile bitterlings of all populations (infected and control groups) were individually collected from netted containers housed in outdoor pools 60 min before being placed in the inner test chamber of the swim tunnel. All individuals were left undisturbed for ∼15 min without active water velocity but a constant flow of oxygenated water to maintain a high dissolved oxygen level inside the tunnel. The fish were then individually subjected to increasing water velocities until they were unable to maintain a steady position against the water flow. Body length (BL), wet weight, final velocity increment and time completed after final velocity increment were recorded once the fish fell to the back of the tunnel, which signalled the end of the trial. U_crit_ values were calculated according to Brett (1965): U_crit_=U_f_ + (t_f_/t_i_U_i_), where U_f_ is the water velocity (BL/s) of the last fully completed increment, t_f_ is the time spent at the last water velocity increment, t_i_ is the period for each completed water velocity increment, and U_i_ is the water velocity increment.

Fish infected with glochidia were sacrificed by an overdose of clove oil and preserved in 4% formaldehyde immediately after the swim tunnel runs to determine their parasite load at the time of the swimming performance test. The number of glochidia attached to the fish was counted under a dissecting stereomicroscope (Olympus SZX7, Japan).

### Overwintering experiment

Control and treatment groups of fish from all populations subjected to the glochidia infection protocol were left undisturbed for 15 days to observe the health condition of juveniles after the infection protocol. The initial number of individuals was 65-80 fish per treatment (Table 1). On 15 September 2023, juvenile fish were measured and randomly allocated to smaller, mesh containers (4 per tank), which were distributed across larger tanks to minimise tank-based biases. All juvenile fish included in the study were fed once daily with an equal ratio of crushed dry flakes and chopped bloodworm.

**Table 1.** Two-week survival of infected and control juvenile bitterling in large fibreglass tubs following experimental infection and χ^2^ test of difference in survival between infected and control groups.

| population | treatment | survival (%) | N | $\chi^2$ | P |
| --- | --- | --- | --- | --- | --- |
| Old | Control | 95.4 | 65 | 6.77 | 0.009 |
|  | Infected | 78.5 | 65 |  |  |
| Intermediate | Control | 93.8 | 65 | 9.25 | 0.002 |
|  | Infected | 72.3 | 65 |  |  |
| Without | Control | 96.4 | 56 | 18.18 | <0.001 |
|  | Infected | 63.8 | 80 |  |  |

On 1 November 2023, we measured the body length of fish in the mesh baskets, which were sealed and moved to a large outdoor pool (5 ×□12 m) dedicated to experimental work. Fish in the baskets were fed with chopped bloodworm, with equal rations among baskets. Water temperature followed a natural seasonal cycle and fluctuated between 4.1□ and 12.0□ during the overwinter period. In spring (20 March 2024), the mesh baskets were recovered from the large pool and transported back to the smaller fibreglass tanks for ease of handling. Fish body length was remeasured to determine growth rates during the overwintering period. On 1 April 2024, fish were sacrificed with an overdose of clove oil and stored in a 4% solution of formaldehyde.

In the laboratory, all fish were measured for their BL to the nearest 1 mm and weighed (W) to the nearest 10 mg. Sex (female, male, or juvenile) was determined by visual assessment of secondary sexual characteristics (e.g., presence of an ovipositor) and confirmed during dissection. Fish were dissected and their gonads extracted, weighed (WG) to the nearest 0.1 mg, and preserved in glycerine. Oocytes were photographed using a stereomicroscope (Nikon SMZ1000, Japan), counted, and measured to the nearest 0.001 mm using LUCIA 5 image analysis software. Absolute fecundity (FA) was estimated gravimetrically as the total number of oocytes per female. Relative fecundity (FR) was expressed as the number of oocytes per 1 g of female body weight.

### Data analysis

Immediate infection effects included estimates of initial glochidia load, swimming capacity and immediate survival. Initial glochidia load was compared among populations using a linear model with population identity as a fixed factor and body length as a covariate. Tukey tests (calculated in the *emmeans* package) were used for pairwise comparisons. Data on swimming capacity, defined as critical swimming ability and calculated per body length unit (Beamish 1978; Jain et al. 1997), was modelled with treatment (infected, control), population identity (without, intermediate, old) and their interaction as fixed factors. An interaction was interpreted as indicating population-specific response to glochidia load. To quantify the immediate effect of glochidia load on fish condition, we tested how the number of glochidia attached to individual fish affects critical swimming ability (per body length), with population identity, and interaction between population and number of glochidia as fixed factors. ANOVA with Type II test was used to estimate the statistical significance of the effects. The *DHARMa* package was used to validate models. Survival between experimental infection and transfer to experimental basket (14 days after infection) was tested using a chi-squared test, as fish from a particular combination of population and treatment were housed in single containers (i.e. six containers in total).

For long-term effects, we considered the growth, overwinter survival, body condition, and reproductive allotment of fish. Growth rates were compared between infected and control fish using measurements of juvenile fish taken on 15 September 2023, 1 November 2023 and 20 March 2024. We compared BL as a function of treatment (infected, control), date (September, November, March), population identity (without, intermediate, old) and their two-way interactions. The initial model included a three-way interaction, but it was not statistically significant (χ^2^ = 0.75, P = 0.685) and hence was dropped from the final model. The main focus of this analysis was on the time by treatment interaction, as it provided information on whether infected and control fish differed in their growth. Fish of each “treatment by population” combination were distributed in four different baskets, and we included basket ID as a random slope to account for the non-independence of fish body length estimates from the same basket. The analysis was conducted in the *glmmTMB* package, using a Gaussian distribution. The *DHARMa* package was used to validate models.

Overwinter survival was tested using binomial GLMM analysis, with the ratio of introduced vs surviving fish as the response variable, and treatment (infected, control) and population (old, middle, without) as fixed factors. Each experimental population and treatment were replicated four times. One experimental basket was damaged, and fish escaped between 1 November and 20 March. We have removed data from that basket from the final dataset. We use survival from 15 September 2023 to 20 March 2024 for the main analysis. The analysis of other date combinations (including 1 November 2023) provided concordant results.

Reproductive allotment was initially measured using a set of reproductive traits. In females, oocytes were counted, their diameter measured, and the GSI was calculated following Wootton (1998) using the formula: GSI = (100 × WG) / W, where WG is gonad weight and W is total body weight. The Fulton condition factor (K) was calculated according to Le Cren (1951) and Ricker (1975) using the equation: K = 100 × W × TL□³, where W is fish weight (g) and TL is total length (cm). Many estimated traits were highly correlated. We used the number of oocytes and GSI (which were not significantly correlated: Pearson correlation r = 0.132, P = 0.264) to formally test the effect of experimental infection on reproductive allotment. Both models contained treatment, population identity and their interaction as fixed effects and BL as a covariate. Data were modelled using a linear model. For the number of oocytes, a Poisson error distribution was also fitted, but the model had lower explanatory power (ΔAIC: 600) than the linear model.

## RESULTS

### Immediate effects

Bitterling populations with longer associations with *S. woodiana* had the lowest initial glochidia load (ANOVA: F_2,31_ = 7.12, P = 0.003, contrast between old and without populations: P = 0.002, contrast between old and intermediate populations: P = 0.056, contrast between intermediate and without populations: P = 0.433) (Fig. 1A). There was a negative association between fish size and glochidia load (F_1,31_ = 4.62, P = 0.040).

**Fig. 1.**
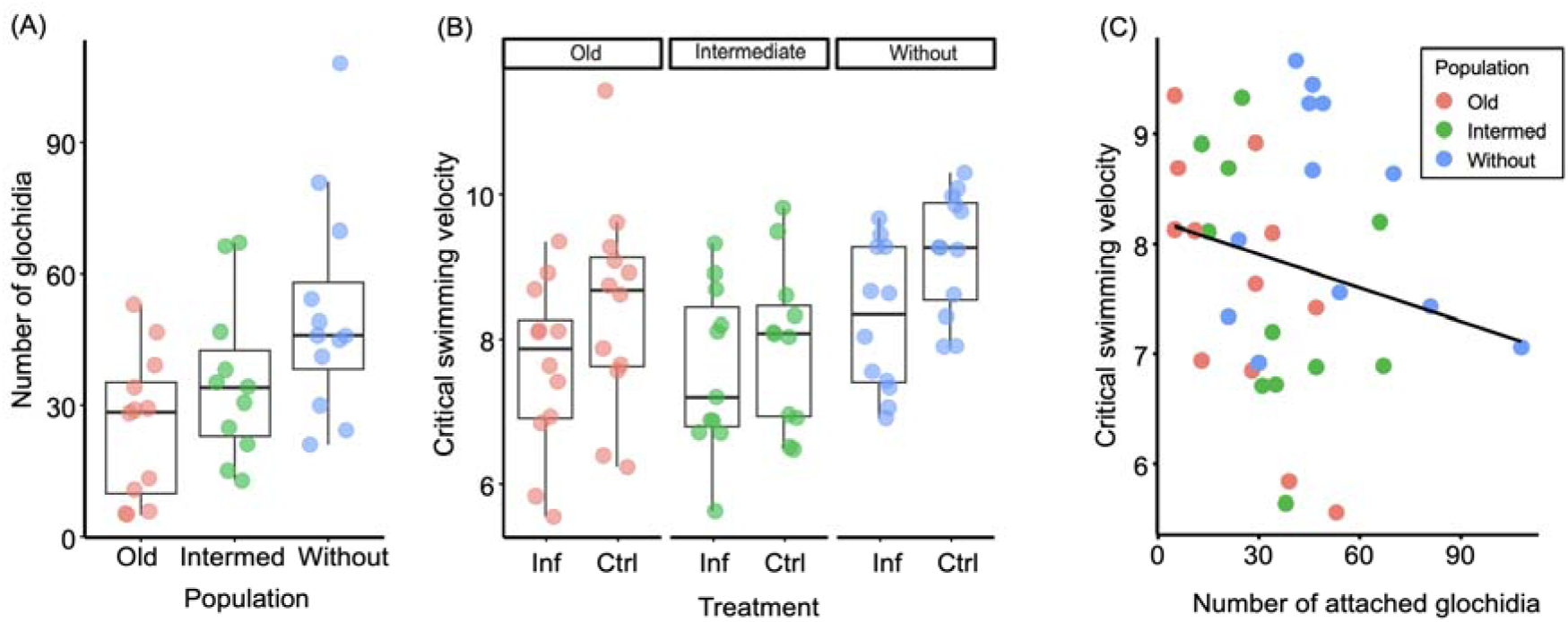
The load of *S. woodiana* after experimental infections (A), critical swimming velocity (B) and their relationship (C) across treatments and three study bitterling populations. Dots represent raw values, boxes denote medians and interquartile range, and whiskers represent non-outlier range. Specific colours denote populations, and the relationship between critical swimming velocity and number of glochidia is visualised using the *lm* plotting function in the *ggplot2* package

Fish with acute glochidia infection had poorer swimming abilities. Experimentally infected fish had a lower swimming capacity than control fish (F_1,64_ = 7.01, P = 0.010) but the response was not population-specific (F_2,64_ = 0.39, P = 0.679) (Fig. 1A). The effect is contingent upon glochidia load severity; within infected fish, higher glochidia load was associated with decreased swimming capacity (F_1,29_ = 6.04, P = 0.020) (Fig. 1B). This analysis confirmed that populations varied in their experimental glochidia load (F_2,29_ = 3.91, P = 0.032), but demonstrates that the effect of glochidia load was not population-specific (interaction: F_2,29_ = 1.53, P = 0.233).

There was significantly increased fish mortality during the initial two-week interval after experimental infections across all three populations. The effect was strongest in the population without any prior coexistence with *S. woodiana* and weakest in the population with an old association with *S. woodiana* (Table 1).

### Long-term effects

There was no decrease in fish overwinter survival after experimental infection. Control fish (85.9 %) and infected fish (88.6 %) did not differ in their survival rate (GLM with binomial error, χ^2^ = 0.13, P = 0.724) (Fig. 2A). There was no difference in growth between infected and control fish (Linear Mixed Model, treatment by time interaction: χ^2^ =0.82, P = 0.366, Figure 3). Infected fish were in better condition than control fish at the end of the experiment (F_1,158_ = 9.37, P = 0.003), with no difference among populations (F_2,158_ = 2.29, P = 0.104, treatment by population interaction: F_2,158_ = 0.26, P = 0.775) (Fig. 2B) and no difference between males, females and juveniles (F_2,158_ = 0.95, P = 0.390). Larger fish had higher values of Fulton’s condition coefficient (F_1,158_ = 10.74, P = 0.001) (Fig. 2D).

**Fig. 2.**
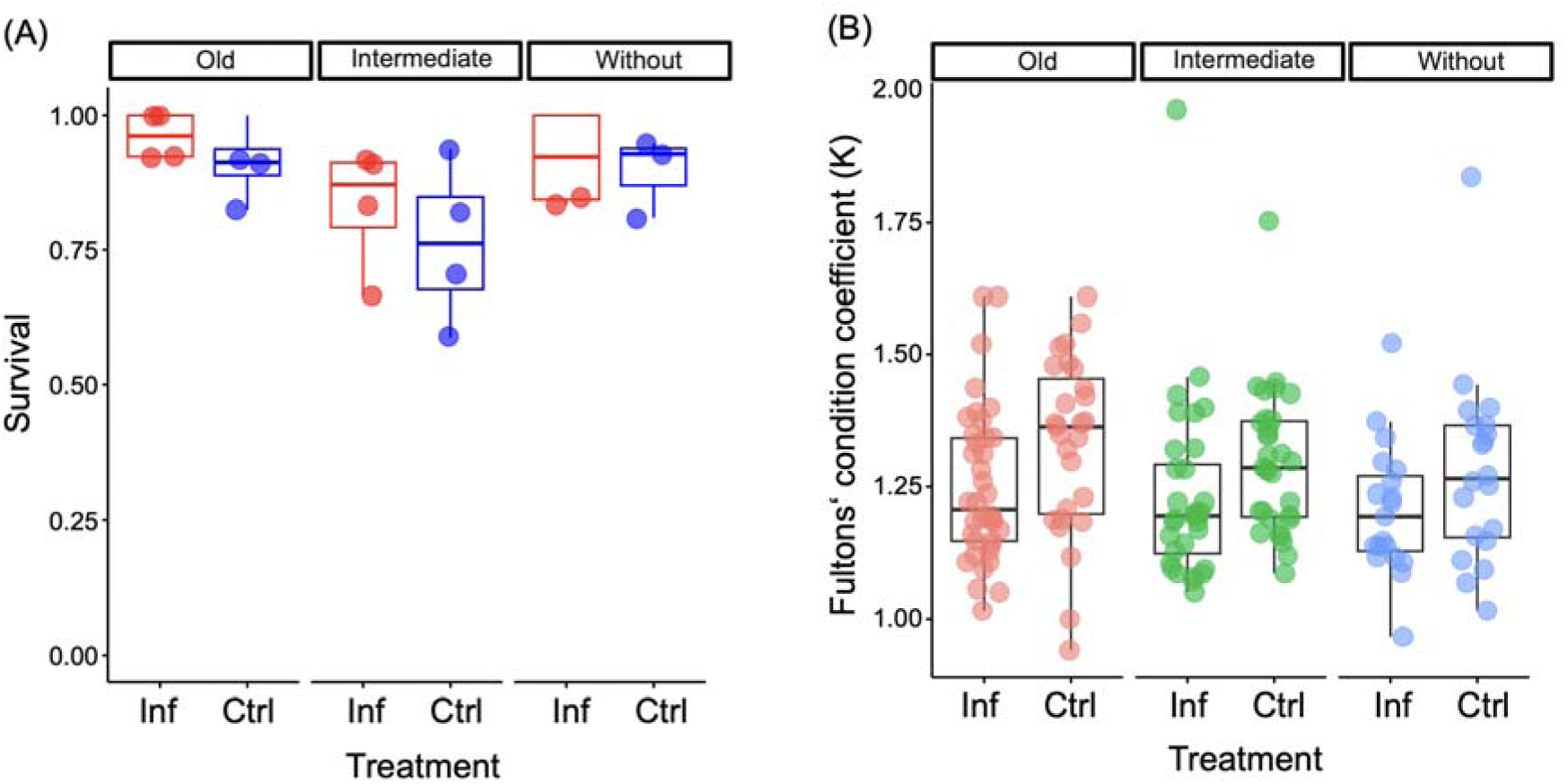
Survival between September 2024 and March 2024 (A) and Fulton’s condition coefficient (B) for every treatment and study bitterling populations combination. Dots represent raw values, boxes denote medians and interquartile range, and whiskers represent non-outlier range

**Fig. 3.**
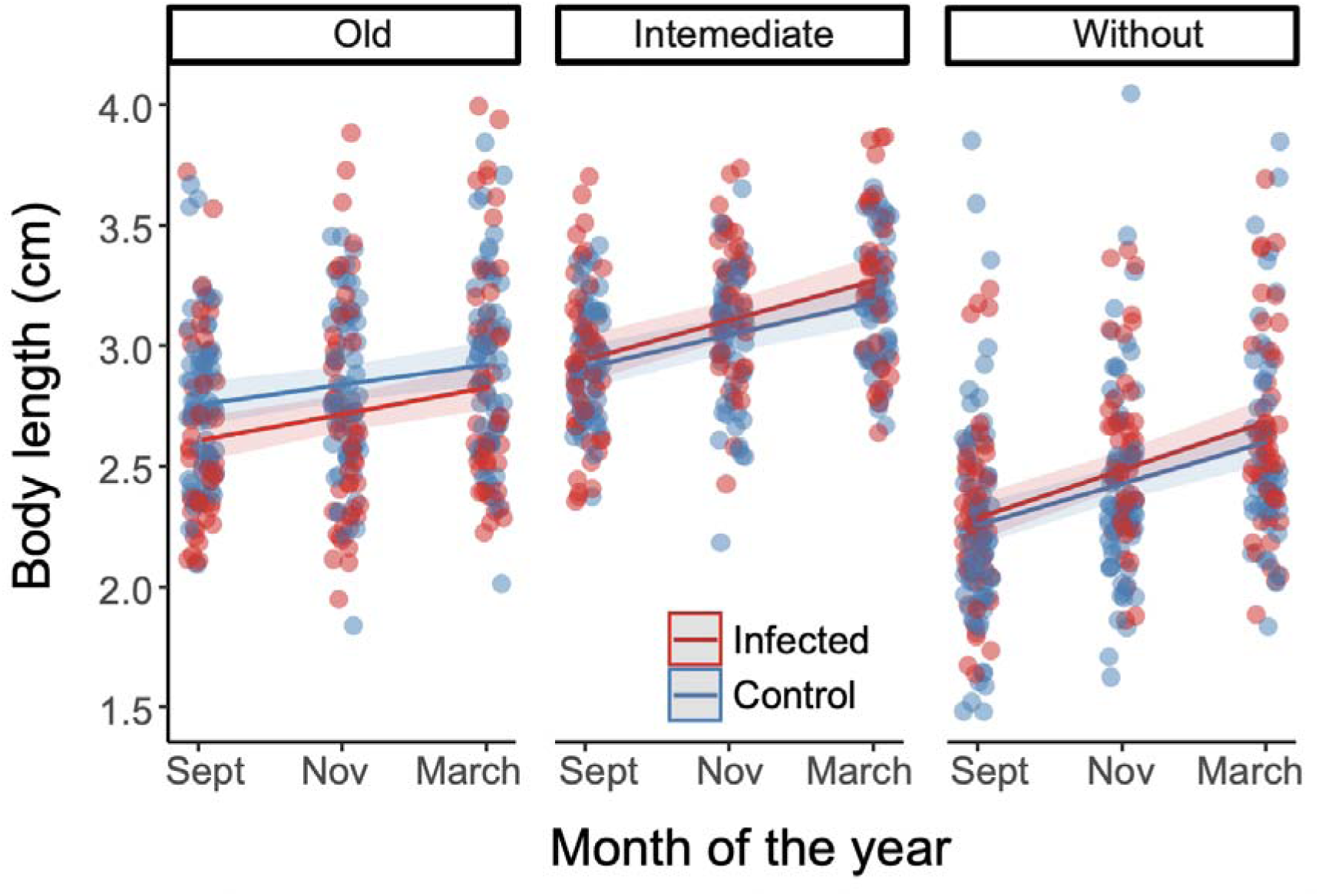
Individual body lengths measured in September 2023, October 2024 and March 2024 in experimentally infected (red) and control (blue) bitterling across three study populations. Model estimated growth and their 95% confidence intervals are shown by lines and shaded areas, respectively. Note that primary interest of the analysis was in growth rates (i.e. whether the growth lines are parallel). Graph was plotted using *sjmodel* function

There was no difference in the number of oocytes between infected and control females (F_1,68_ = 0.71, P = 0.402) nor population-specific cost of infection (interaction effect: F_2,68_ = 0.73, P = 0.485). Fish from different populations varied in their oocyte count (F_2,68_ = 7.96, P =0.001) (Fig. 4A). Larger females tended to have more oocytes (F_1,68_ = 3.85, P = 0.054) (Fig. 4B). Similarly, there was no effect of experimental infection on female GSI (F_1,68_ = 1.54, P = 0.220, population by treatment interaction: F_2,68_ = 1.31, P = 0.277) (Fig. 4C). The GSI did not vary among populations (F_2,68_ = 0.40, P =0.670) but was strongly positively related to female BL (F_1,68_ = 45.06, P < 0.001) (Fig. 4D). Overall, infected and control males did not differ in their GSI (F_1,67_ = 0.07, P = 0.787) (Fig. 5A, B). There was no difference in male GSI among populations (F_2,67_ = 0.41, P = 0.669). However, there was a population–specific response to infection in male reproductive allotment (F_2,67_ = 7.86, P = 0.001); infected males had lower GSI than control males in the old-association population but not in the other two populations (Fig. 4A).

**Fig. 4.**
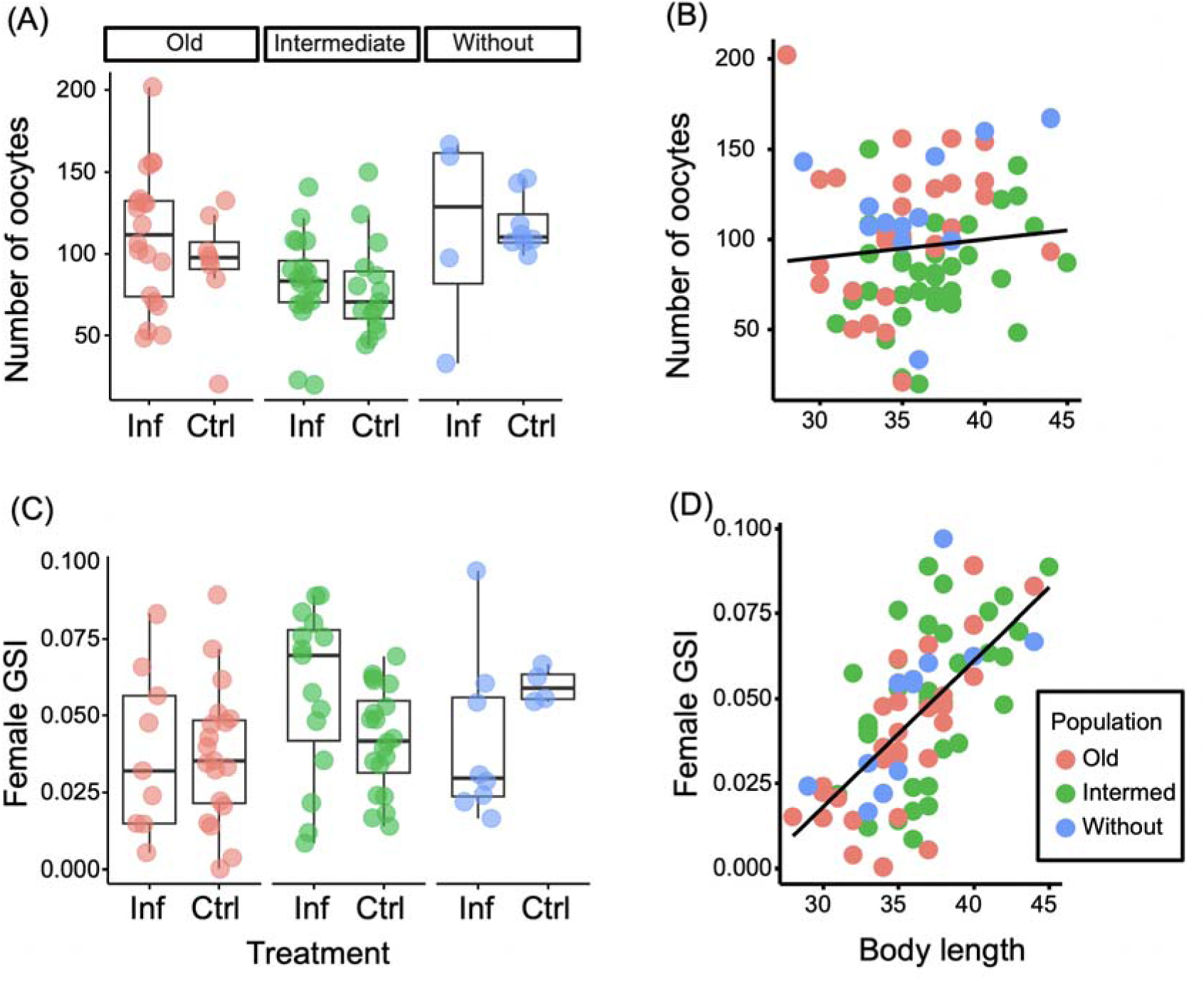
Female reproductive traits across treatments and three study bitterling populations. The number of oocytes (A, B) and female gonadosomatic index – GSI (C, D) are visualised as boxplots for each treatment by population combination (A, C) and in relation to individual body length (B, D). Dots represent raw values, boxes denote medians and interquartile range, and whiskers represent non-outlier range. Specific colours denote populations, and the relationship between reproductive parameters and body length is visualised using the *lm* plotting function in *the ggplot2* package

**Fig. 5.**
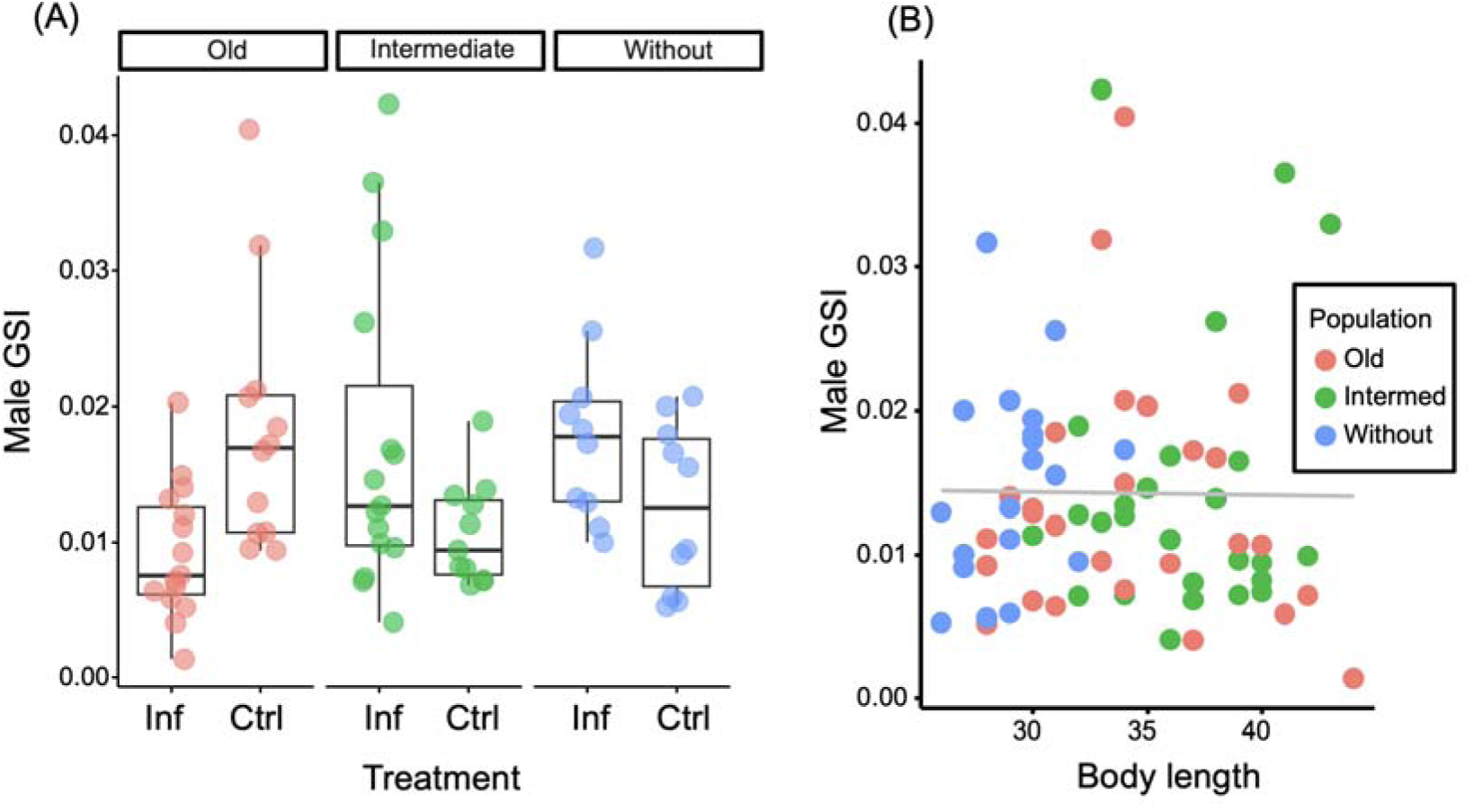
Male gonadosomatic index (GSI) across treatments and study bitterling populations (A) and in relation to individual body length (B). Dots represent raw values, boxes denote medians and interquartile range, and whiskers represent non-outlier range. Specific colours denote populations, and the non-significant relationship between male GSI and body length is visualised by a grey line using *the lm* plotting function in the *ggplot2* package

## DISCUSSION

We studied the immediate and long-term costs of infection by parasitic larvae of the invasive mussel *Sinanodonta woodiana* on host fish *Rhodeus amarus* across a temporal gradient of the history of their interaction. In this interaction, glochidia of *S. woodiana* readily use *R. amarus* as a host, with an infection period of 1-2 weeks before detachment and metamorphosis (Douda et al. 2012, 2017b, Anil et al. 2024). Here, we demonstrate that the fish population with the longest evolutionary coexistence with the invasive parasite was least infected. We further demonstrate significant immediate effects, but no long-term effects, of experimental infection of *R. amarus* by *S. woodiana*. Specifically, parasitised fish suffered reduced swimming performance during the initial glochidia attachment period and elevated mortality during the first two weeks after experimental infection. In the infected fish that survived the initial period (64-79 %), no decrease in their long-term (overwinter) survival or decrease in somatic condition and reproductive investment in the subsequent spring was detected. Traits such as growth rate, GSI, and oocyte number remained unaffected by glochidia infection. Bitterling populations were consistent in their response to glochidia load across all measured long-term parameters, suggesting that the major fitness costs of parasitism are concentrated in the initial period following infection, possibly due to acute tissue damage and physiological stress responses, as reported in other similar systems (Douda et al. 2017a). We discuss potential explanations for the observed survival differences across populations and implications for rapid parasite-mediated selection.

The swimming capacity of infected fish was significantly reduced. The effect was load-dependent, with individual fish carrying higher glochidial loads exhibiting lower swimming performance. As predicted, increased glochidia attachment hindered fish from reaching optimal swimming speeds. This impairment likely results from glochidia attaching to fins and gills, thereby disrupting fin movement and reducing oxygen uptake. Similar effects have been reported in other host–parasite systems, such as reduced swimming speed in the coral reef fish *Scolopsis bilineatus* infected by an ectoparasitic isopod (Binning et al. 2012). Swimming performance in juvenile fish is vital for survival, influencing both foraging and predator evasion. In natural settings, reduced swimming ability decreases foraging activity (Nunn et al. 2012) and ability to evade predators, with empirical evidence from various taxa including larval fish, frogs and damselflies (McPeek 1996; Wilson et al. 2005; Masuda et al. 2006). Larval and juvenile jack mackerel use burst swimming to evade jellyfish (Masuda et al. 2006), and damselfly larvae have repeatedly evolved enhanced swimming abilities in response to predatory dragonflies (McPeek 1996). Given that swimming constitutes a substantial portion of the total bioenergetic budget in small fish (Ruzicka and Gallager 2006; Brett and Groves 1979), the combination of reduced oxygen uptake and impaired locomotion caused by glochidia parasitism likely exerts additive stress on juvenile bitterling.

Regarding the glochidia load of *S. woodiana*, we observed significant population-level differences in glochidia attachment among bitterling fish. Consistent with our prediction on the evolution of increased resistance, bitterling populations with longer coexistence with *S. woodiana* suffered lower glochidia loads. During the first four days post-infection, individuals from the population with the oldest association with *S. woodiana* carried 30% and 51% fewer glochidia than those from intermediate and naïve populations, respectively. This pattern indicates rapid evolution leading to higher resistance to *S. woodiana* glochidia. Although the mechanisms behind this resistance remain to be confirmed (e.g. behavioural avoidance, mucus shedding, or innate immune activation) (Donrovich et al. 2017; Rock et al. 2025), the observed pattern strongly suggests the action of parasite-mediated selection. Rapid coevolution often shapes trajectories of host–parasite dynamics (Calcagno et al. 2010). As parasitic load is closely linked to host survival, with lower loads generally reducing the physiological cost of infection (Hollanders et al. 2022), our findings highlight the evolution of adaptive host responses likely driven by selective pressure from parasitism.

The immediate survival of juvenile bitterling in the infected groups was consistently lower than in the control groups. Survival rates were 17%, 22% and 33% lower in infected groups in old, intermediate, and no association with *S. woodiana*, respectively (Table 1), reinforcing the possibility of rapid evolutionary response of the bitterling to *S. woodiana* invasion. Host survival is in the interest of the parasite, supporting its development. However, range expansions of parasites into new regions sometimes cause coevolutionary mismatch. For example, a monogenean parasite *Gyrodactylus salaris* caused substantial mortalities of East Atlantic populations of the Atlantic salmon (*Salmo salar*) upon its introduction to the region, despite having no major impact on the salmon populations in the Baltic, where *G. salaris* is likely native and has coevolved with salmon (Bakke et al. 2002). Similar mismatches have been reported in other systems, including increased mortality in bumblebees exposed to non-native parasites (Meeus et al. 2011) and European eels infected by a non-native helminth (Taraszewski 2006). Adaptations to non-native parasites, with consequent host population recovery, are reported for anurans which suffered severe fungal infections (Hollanders et al. 2022), Trinidadian guppies parasitised by a nematode (Rogowski et al. 2020) and multiple rabbit populations decimated by myxovirus (Alves et al. 2019). While these cases exemplify pathogens with strong selection for resistance or tolerance, our results highlight that decades of coexistence with a new parasite may trigger an evolutionary response in less virulent cases.

In contrast to clear short-term effects, we found no significant long-term effects of glochidia parasitism on juvenile bitterling. There was no effect on overwinter survival and no differences in growth rate, condition factor, or reproductive allotment after overwintering. These results did not correspond with our predictions of long-term negative impacts of glochidia parasitism. There was also no evidence of population-specific differences, except for a reduced male GSI in the infected fish group for the old association population. There are two main explanations for this lack of an effect. First, coevolution between bitterling and glochidia of native mussels may have shielded the bitterling from the effect of *S. woodiana* glochidia parasitism. However, resistance rather than tolerance of parasitism was apparent for immediate effects (i.e. lower glochidia load along the history of association), and weak or inconsistent correspondence between susceptibility to native versus invasive glochidia (Douda et al. 2017b; Rouchet et al. 2017). Second, higher immediate mortality in the bitterling populations with intermediate and no history of association with *S. woodiana* may have eliminated weaker individuals, masking any longer-term differences through a process known as selective disappearance. This non-random mortality, commonly observed in wild populations, can shift trait distributions through the elimination of individuals less adapted to their environment. For example, under this scenario, lighter ungulates experience higher mortality across life stages (Nussey et al. 2010), and great tits with shorter telomeres are selectively lost in urban populations, resulting in populations with longer telomeres (Salmon et al. 2016). Similar dynamics may have influenced trait composition in our bitterling populations, potentially concealing the long-term costs of parasitism. We suggest that selective disappearance is a more likely explanation of our experimental results, and it implies a strong evolutionary response to the novel parasite.

Our results show the immediate impact on survival and swimming capacity of *R. amarus* in response to glochidia parasitism by *S. woodiana*, but no significant long-term physiological or reproductive cost. While we performed our experiment during peak glochidia release from maternal mussels, glochidia production in *S. woodiana* also occurs at other times of the year (Douda et al. 2012) and, in some *S. woodiana* populations in Europe, even continuously (Labecka and Domagała 2018). Consequently, the immediate impacts may extend beyond those recorded in this study. *S. woodiana* is capable of taking advantage of evolutionarily naïve hosts in the areas of range expansion. As a generalist host fish parasite, *S. woodiana* threatens not just host fish species but also other native mussels, including rare and endangered species (Donrovich et al. 2017; Huber and Geist 2019; Urbańska et al. 2021; Lewisch et al. 2023; Halabowski et al. 2024). Our study adds a temporal dimension to this phenomenon, suggesting that even after decades of coexistence, host populations may unergo measurable evolutionary responses. The observed population-level differences in response to *S. woodiana* parasitism are consistent with parasite-mediated selection, although further work is needed to confirm heritability and the genetic basis of these traits. Similar rapid adaptive responses to novel interactions have been observed across diverse taxa. For instance, the seed-feeding soapberry bug (*Jadera haematoloma*) evolved morphological changes within a few decades after switching to an introduced host plant (*Cardiospermum halicacabum*) (Carroll et al. 2001). Similarly, the invasion of *Anolis sagrei* in Florida prompted rapid adaptive shifts in perch height and limb morphology in native *Anolis carolinensis* lizards (Stuart et al. 2014).

Biological invasions present outstanding models to study ecological and evolutionary mechanisms in nature (Feis et al. 2016, 2019) and the consequences of anthropogenic exchange of species between environments (Mooney and Cleland 2001). Ultimately, the interplay between parasite-mediated selection and population-level host responses may drive the evolution of resistance, as reflected by reduced glochidia load and higher survival in populations with a longer history of coexistence. Our findings suggest that novel host– parasite interactions established during biological invasions can trigger measurable adaptive responses over relatively short ecological timescales and, in consequence, rapidly change the impact of non-native species on invaded ecosystems. It highlights the importance of considering evolutionary responses and spatiotemporal variation in the associations between invading and native species during the risk assessment and management of biological invasions. Future research disentangling the contributions of genetic adaptation, phenotypic plasticity, and selective disappearance will be essential to fully understand the evolutionary trajectory and stability of such associations.

## Acknowledgements

We wish to express our sincerest gratitude to Dr Anna Maria Łabęcka for providing data on *S. woodiana* localities and other essential information for the selection of study sites. All procedures were approved by the Local Ethics Committee (19/ŁB264/2023) and were carried out in accordance with permission from the General and Regional Directorates of Environmental Protection: General (DZP-WG.6401.111.2023.ASZ.2); Lodz (WPN.6401.325.2022.BWO.3, WPN.672.3.2022.AGr); Cracow (OP.6401.193.2023.GZ, OP.672.38.2022.GZ, OP.6401.67.2023.GZ); Katowice (WPN.672.30.2022.MS1, WPN.6401.252.2023.DT); Poznań (WST.6401.229.2023.MT.2, WST.672.14.2022.MK); Warsaw (WSTR.6401.26.2003.MK.2, WPN-I.672.7.2022.KZ.3, WPN- I.6401.72.2023.MK.2).

## Author Contributions

A.N.A. and D.H.: conceptualisation, methodology, investigation, writing – original draft, writing – review and editing; K.P., G.Z., J.G., K.D. and C.S.: investigation, writing – review and editing; M.R.: conceptualisation, methodology, investigation, visualisation, formal analysis, writing – original draft, writing – review and editing, supervision.

## Data availability

All data generated or analysed during this study were uploaded to the Figshare repository (https://doi.org/10.6084/m9.figshare.29617205).

